# Reduced intra- and inter-individual diversity of semantic representations in the brains of schizophrenia patients

**DOI:** 10.1101/2020.06.03.132928

**Authors:** Satoshi Nishida, Yukiko Matsumoto, Naganobu Yoshikawa, Shuraku Son, Akio Murakami, Ryusuke Hayashi, Shinji Nishimoto, Hidehiko Takahashi

## Abstract

Schizophrenia patients often manifest semantic processing deficits. It has been proposed that these deficits stem from disorganized semantic representations in the brain. However, no study has yet examined the neural correlates of semantic disorganization by directly evaluating semantic representations in the brain. We used voxelwise modeling on functional magnetic resonance imaging signals to evaluate the semantic representations associated with several thousand words in individual patient brains. We then compared the structural properties of semantic representations to those in healthy controls. The variability of semantic representations was smaller both within individual patients and across patients compared to controls. Surrogate data analysis suggests that the observed reduction in representational variability is associated with disorganization of categorical information. To our knowledge, these findings provide the first evidence for sematic disorganization in schizophrenia at the level of brain representations.

## Introduction

Patients with schizophrenia frequently manifest semantic processing impairments^1–3^, which may contribute to the formal thought disorder that is a hallmark of this disease^3–6^. It has been proposed that semantic processing impairments in schizophrenia stem from the disorganization of semantic storage in the brain^7,8^ as evidenced by behavioral tests, such as verbal fluency^9–14^. However, no study has yet examined the neural correlates of semantic disorganization in schizophrenia by directly evaluating semantic representations in patient brains.

Recent neuroimaging studies have utilized voxelwise modeling to evaluate the organizational structure of semantic representations in individual brains^15–18^. For example, Huth et al. (2012) constructed a voxelwise model that predicts brain activity patterns during the viewing of natural movie clips from object/action word labels for each scene^18^. These analysis yielded representational spaces for thousands of semantic entities, such as “man”, “dog”, and “car”, in individual brains. As an extension of this work, several studies have attempted to model semantic representations in the brain more efficiently using word vector spaces^15,16,19–21^. Word vector spaces learned from text corpora using natural language processing algorithms (e.g., word2vec^22^) can effectively capture the sematic relationships among thousands of words. Accordingly, voxelwise modeling based on a word vector space can comprehensively evaluate the semantic representations associated with thousands of words in the brain^15–17^.

We speculated that such semantic representations quantified by voxelwise modeling based on a word vector space would be disorganized in schizophrenia patients compared to healthy controls. To test this hypothesis, we constructed voxelwise models to predict brain responses of individual brains to the semantic contents in movie scenes (Fig. 1a). Brain response to natural movie scenes were collected from both patient and control groups using functional magnetic resonance imaging (fMRI) and the semantic contents in each scene were manually annotated in the form of short descriptions by independent observers. The scene descriptions were then transformed into scene vectors through a word vector space learned in advance from large-scale text data using the word2vec algorithm^22^. Drawing on the evoked responses and corresponding scene vectors for each individual, we trained a voxelwise model using linear regression. Then, the regression weights were used to evaluate the word representations in individual brains (brain word representations) from those in the word vector space (Fig. 1b). These brain word representations correspond to predicted brain responses for semantic information associated with individual words. Finally, the structural characteristics of brain word representations were compared between schizophrenia patients and healthy controls.

**Fig. 1.**
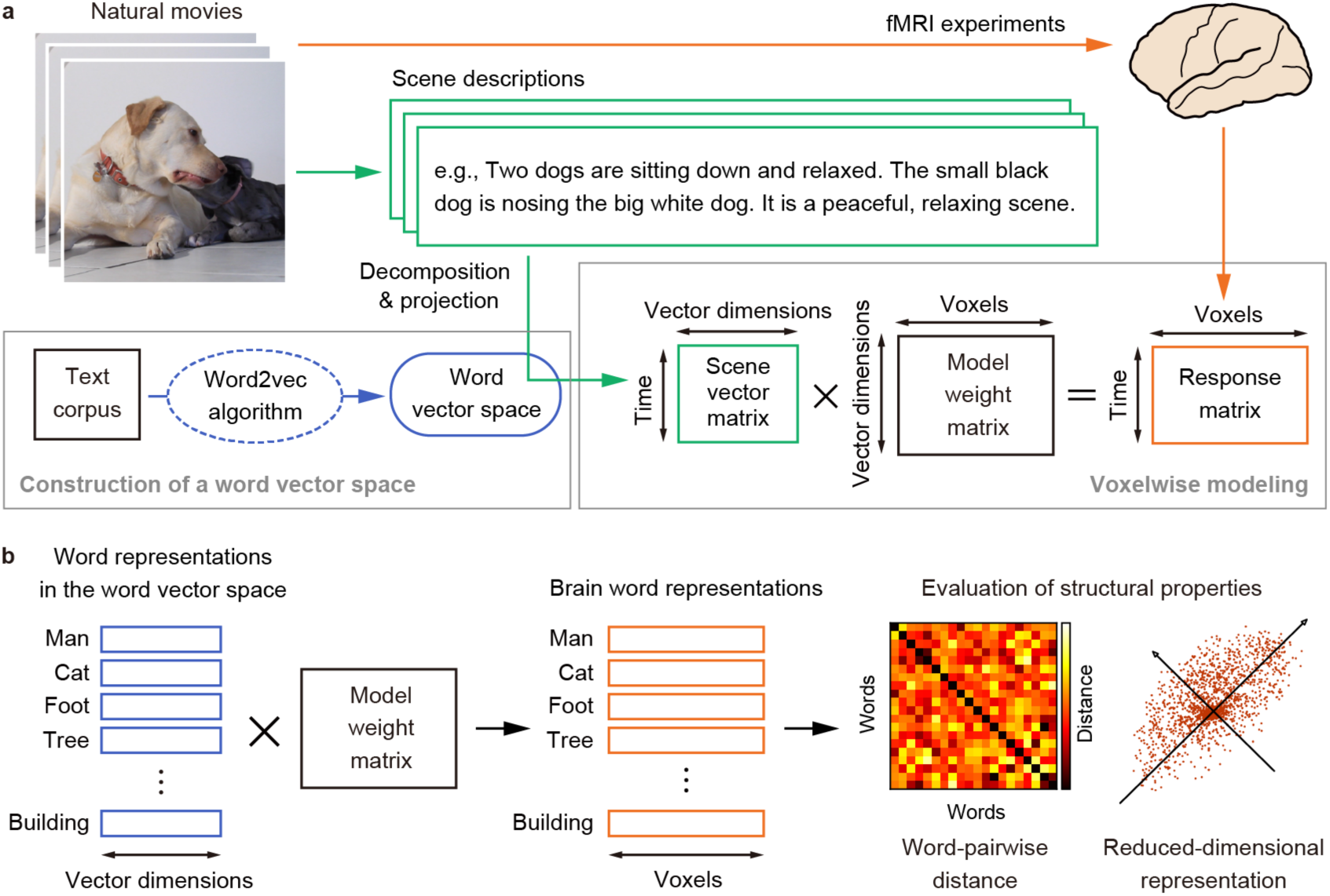
Modeling of semantic representations in the brain. **a**, Voxelwise modeling based on a word vector space. The model predicts individual brain responses to natural movie scenes by a weighted linear summation of the vector representations of semantic contents in each scene (scene vectors). The scene vectors were obtained by transforming manually annotated movie scene descriptions through a word vector space constructed in advance from a text corpus using a language processing algorithm (word2vec^22^). **b**, Data analysis scheme for evaluating the structural properties of semantic representations in the brain. These semantic representations were evaluated by predicted brain responses to the semantic information associated with individual words (brain word representations). The brain word representations were computed by multiplying original word vectors in the word vector space by the model weights. Then, the structural properties of brain word representations in individual brains were examined using word-pairwise distance and reduced-dimensional representation.

## Results

### Lower intra-individual representational variability in schizophrenia

We first investigated the structural properties of brain word representations within subjects using two separate measures: *word-pairwise distance*, which reflects how the dissimilarity between each pair of semantic representations is structured in the brain, and *reduced-dimensional representation*, which reflects how semantic representations are distributed in particular dimensions of representational space after dimensional reduction by principal component analysis (PCA). The word-pairwise distance was measured using the pairwise correlation distance between all possible pairs of brain word presentations. The mean word-pairwise distance did not differ between groups, but the distribution was narrower (i.e., the variance was significantly lower) in patients than controls (Fig. 2a–c; Mann-Whitney U test, P < 0.001). Next, the variability in the reduced-dimensional representation was measured by the explained variance ratio of each PC. The explained variance ratio of the first PC was significantly smaller in patients (Fig. 2d; Mann-Whitney U test, P < 0.005), while the explained variance ratios of the other PCs did not differ (P > 0.05). Together, these analyses indicate that schizophrenia patients have lower intra-individual variability in semantic representations than healthy controls.

**Fig. 2.**
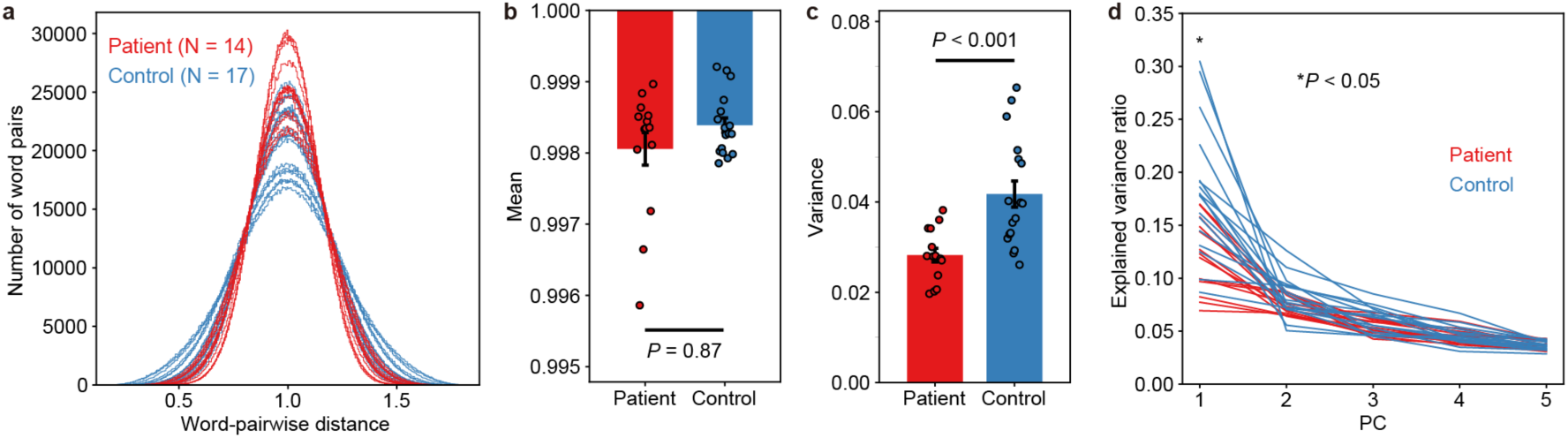
Reduced intra-individual variability in brain word representations among schizophrenia patients. **a**, Distributions of word-pairwise distances computed as the correlation distances between all possible pairs of 2000 brain word representations. Red and blue traces depict the distributions for individual patients and controls, respectively. **b**,**c**, Comparisons of word-pairwise distance distribution means (**b)** and variances (**c)** between patients and controls. Bars depict the means of the statistical parameters and error bars indicate the standard error of the mean. Each circle depicts an individual subject. There was a significant group difference in variance (P < 0.001) but not mean (P = 0.87). **d**, Variance ratio explained by each principle component (PC). The PCs were computed using PCA on brain word representations for each subject. Red and blue traces depict the explained variance for individual patients and controls. There was a significant group difference for the 1st PC (P < 0.005) but not for the other PCs (P > 0.05)

This lower intra-individual variability in the patient group is not due to a difference in voxelwise model accuracy (i.e., how accurately the model explains the semantic representations in individual brains) as this did not differ between groups (Supplementary Fig. 1). Moreover, the distribution of voxels, selected for calculating brain word representations, across cortical regions in individual brains did not differ between groups (Supplementary Fig. 2 and Supplementary Table 1).

This reduced variability in patients, however, does not mean that semantic representations are randomly distributed, but only that representations are not as well organized as in controls. By randomly shuffling the dimensions of brain word representations (i.e., voxels) in individual brains (see Methods), we can destroy the arrangement while holding the variance constant in each dimension. In this case, the variance of word-pairwise distances and the explained variance ratio of the first PC were drastically reduced in both groups (Supplementary Fig. 3), indicating that the semantic representations of patients are structured but not as precisely as those of controls.

### Lower inter-individual representational variability in schizophrenia

We next investigated the inter-individual similarity of semantic representational organization by calculating Pearson’s correlations between individual word-pairwise distances and reduced-dimensional forms of brain word representations for all subject pairs within groups. Pearson’s correlations between word-pairwise distances were significantly larger among patients than among controls (Fig. 3a; Mann-Whitney U test, P < 10^−9^). Similarly, the subject-pairwise correlations of the first PC (which most strongly reflects differences in brain word representations among subjects; Fig. 2d) were significantly larger among patients than controls (Fig. 3b; Mann-Whitney U test, P < 10^−7^), indicating smaller variability in reduced-dimensional brain word representations across patients. Together, these results indicate that patients have lower inter-individual variability in semantic representations than controls.

**Fig. 3.**
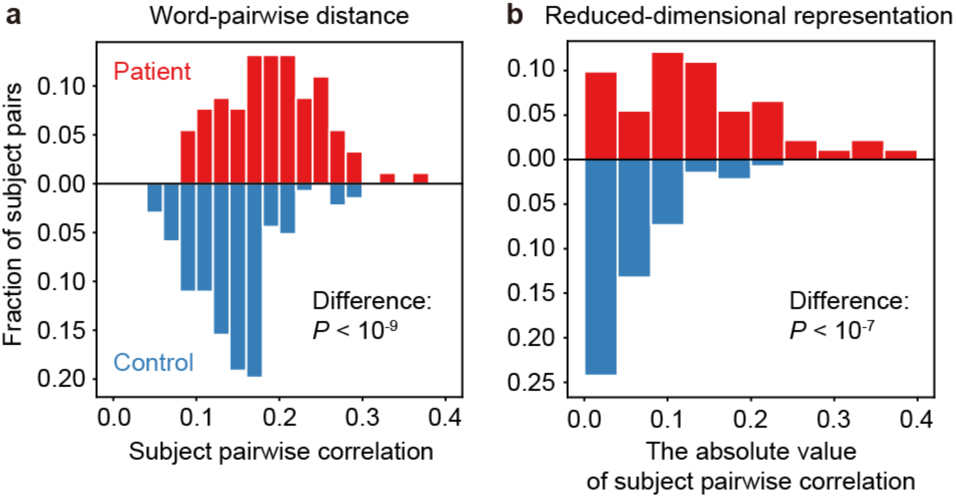
Reduced inter-individual variability in brain word representations among schizophrenia patients. **a**, Distribution of inter-individual similarity in word-pairwise distances. Similarity was evaluated by Pearson’s correlations between word-pairwise distances for each subject pair within groups (subject-pairwise correlations). The difference between groups was significant (P < 10^−9^) **b**, Distribution of inter-individual similarity in reduced-dimensional representations. Similarity was evaluated by the absolute values of Pearson’s correlations between the first PC of brain word representations for each subject pair within groups. The difference between groups was significant (P < 10^−7^)

It is noteworthy that the structural properties after group averaging did not show a clear distinction between patients and controls (Supplementary Figure 4). This indicates that the variability of structural properties at the individual level differs between groups.

### Reduction of representational variability within semantic categories among schizophrenia patients

Semantic representations associated with words exhibit categorical structure (organization) within the continuous representational space of the brain. In other words, groups of semantically related word representations (e.g., “human”, “animal”, and “vehicle”) form clusters in the representational space^16,18,23^. To evaluate if representational variability in schizophrenia is also reduced in these categorical structures, we examined intra- and inter-individual variability of brain word representations using six separate semantic categories, “human”, “animal”, “natural thing”, “body part”, “building”, and “vehicle”, each including 25 unique words.

First, we evaluated the intra-individual variability in word-pairwise distances of brain word representations within each semantic category (within-category distances) for each subject and compared results between groups. The within-category variance averaged over all six categories was significantly lower for patients than controls (Fig. 4a; Mann-Whitney U test, P < 0.005; see Supplementary Fig. 5 for each category). However, intra-individual variability of within-category brain word representations decreased greatly after dimensionality shuffling (Supplementary Fig. 6), suggesting that these brain word representations are not randomly distributed in patients (i.e., there is still categorical structure), but rather contain some fine-scale structures within each category. Thus, the fine-scale semantic organization of brain word representation appears weaker in patients than controls (Fig. 4a).

**Fig. 4.**
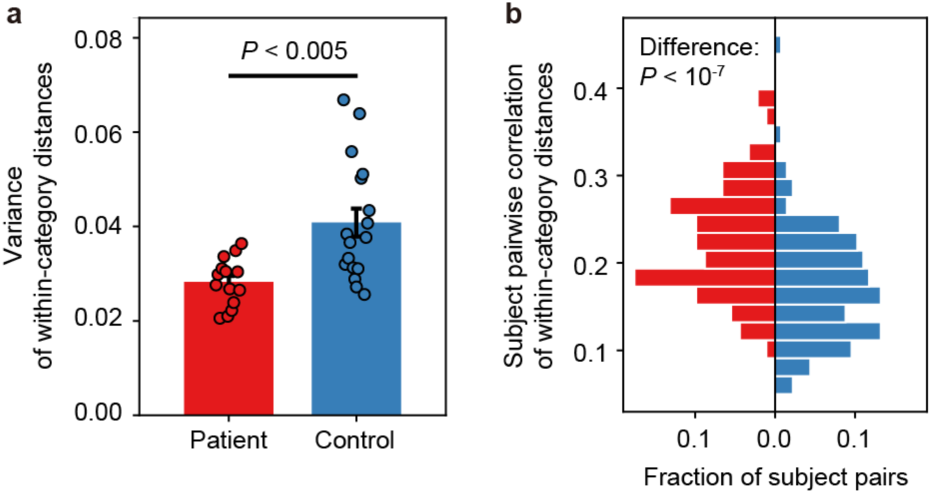
Reduced variability in semantically categorized brain word representations among schizophrenia patients. **a**, The intra-individual variability of brain word representations within word categories. Each category contains 25 unique semantically related words. For each subject, word-pairwise distances within each category (within-category distances) were computed and the variance averaged over all categories and compared between groups. The subject-group difference was significant (P < 0.005). Same conventions as Fig. 2c, but with the vertical axis showing the variance of within-category distances. **b**, Inter-individual variability of brain word representations within word categories. Subject-pairwise correlations were computed within groups using the within-category word-pairwise distances of brain word representations. The red and blue bars show the subject-pairwise correlations for patients and controls, respectively. The subject-group difference was significant for both distances (P < 10^−7^).

Next, we evaluated the inter-individual variability of brain word representations using the same word categories by computing subject-pairwise Pearson’s correlations of within-category word-pairwise distances. Again, these correlations were significantly higher among patients than controls (Fig. 4b; Mann-Whitney U test, P < 10^−7^; see Supplementary Fig. 7 for each category), indicating that the inter-individual variability of brain word representations is lower within semantic categories among schizophrenia patients.

The categories used in these analyses consisted of arbitrarily selected word sets from the vocabulary. However, the specific words chosen were not critical to the results as evidenced by categorical structures in brain word representations using the x-means unsupervised clustering method^24^ (see Methods for details). The x-means clustering is an extension of the k-means clustering and automatically estimates not only cluster structures in distributed data but also the optimal number of clusters for the data. The mean optimal number of clusters was 6.5 for patients and 7.8 for controls, indicating that brain word representations have categorical structures in both groups. Despite the smaller optimal number of clusters for patients than controls (Supplementary Fig. 8a), the intra-individual variability of within-category (within-cluster) word-pairwise distances was lower for patients (Supplementary Fig. 8b), consistent with the results when categories consisted of arbitrary sets of words (Fig. 4a).

### Disruption of within-category structures underlying reduction of representational variability

Our results reveal that the intra-individual representational variability is lower in schizophrenia patients than controls not only for words in whole (Fig. 2) but also for words within specific semantic categories (Fig. 4a). To identify specific differences in representational structure between groups underlying this reduced variability, we performed computer simulations in which the structures of brain word representations were manipulated (Fig. 5) as detailed below.

**Fig. 5.**
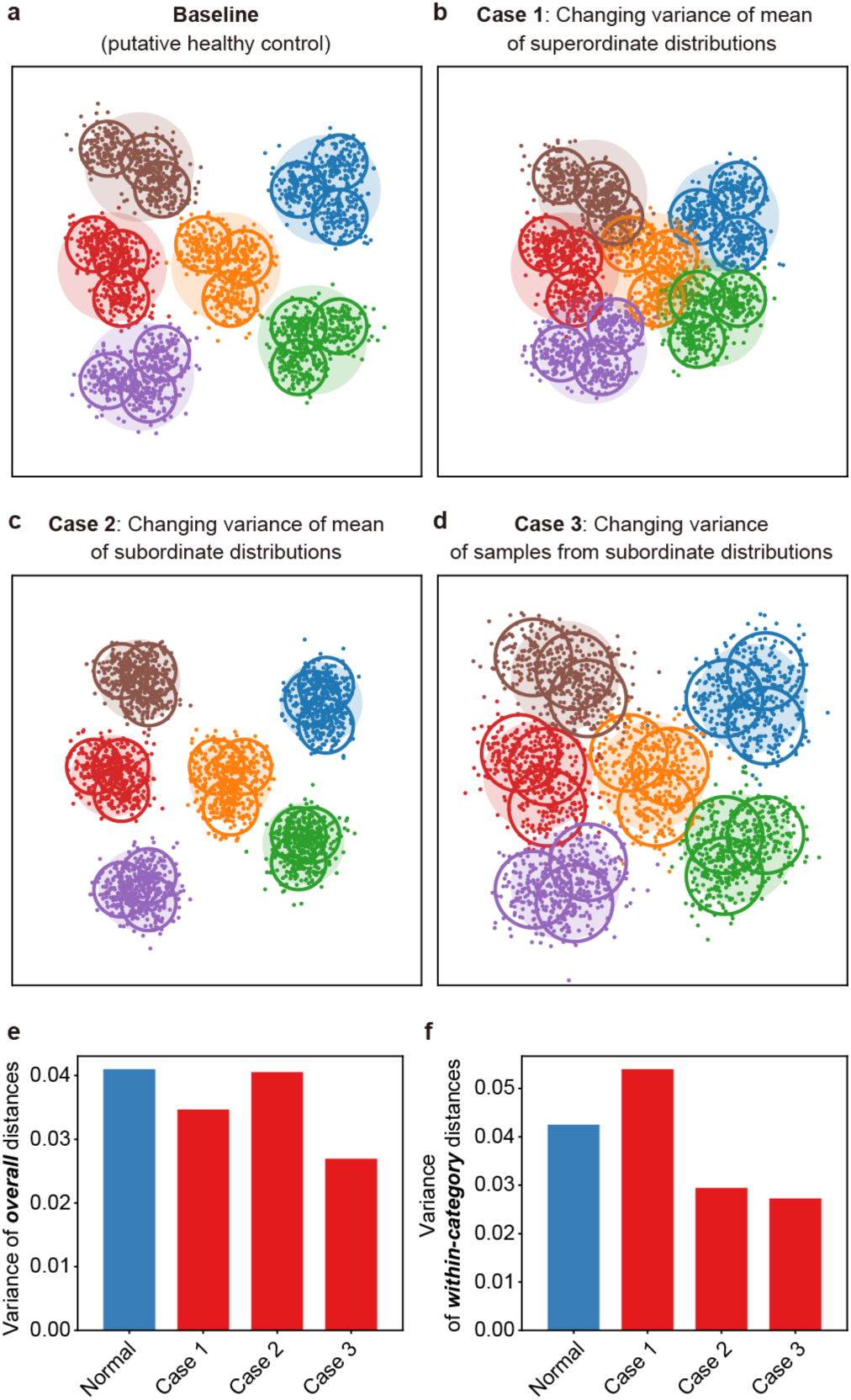
Computational simulations explaining reduced intra-individual representational variability in patients. **a–d**, Illustrations of simulated structural changes in brain word representations. In these simulations, categorical structures were mimicked by six superordinate Gaussian distributions (large filled circles) and fine-scale structure within each category by three subordinate Gaussian distributions (mid-size open circles), the mean parameters of which were drawn from each superordinate distribution. Samples corresponding to brain word representations were drawn from each subordinate distribution (small dots). Different colors indicate different categories. We simulated structural changes in possible cases of patients compared to putative controls (**a**) by manipulating the variance of superordinate distribution means (**b**; case 1), the variance of subordinate distribution means (**c**; case 2), or the variance of samples from subordinate distributions (**d**; case 3). We then examined the effects of these parametric changes on the variability of word-pairwise distances. Note that although the illustration is shown in two-dimensional space for visualization, the computer simulation was conducted on a much higher-dimensional space. **e–f**, Variance in overall (**e**) and within-category (**f**) pairwise distances of brain word representations in the simulations of putative controls (blue bars) and the structural changes for all three cases in patients (red bars). Only in case 3 do the variances of overall and within-category pairwise distances decrease relative to the putative controls and yield a reduction pattern consistent with fMRI data (Figs. 2 and 4**a**; see also Supplementary Fig. 9).

We have so far demonstrated robust structural organization of brain word representations in both patients and controls, albeit stronger in the control group (Supplementary Fig. 3). Such structures contain multiple levels or scales, and we were able to distinguish at least two levels. The first is categorical structure, which corresponds to the separation of representations into individual semantic categories (Supplementary Fig. 8), in accord with previous findings^16,18,23^. The second is fine-scale structure within individual categories (Supplementary Fig. 6).

We speculated that changes in categorical and/or within-category structure among patients would be associated with the observed reduction in intra-individual variability and performed computer simulations to test this hypothesis. We simulated categorical and within-category structures using nested Gaussian distributions (Fig. 5a). The categorical structures were characterized by six superordinate Gaussian distributions, each controlling three subordinate Gaussian distributions. The means of the subordinate distributions were drawn from the corresponding superordinate distribution. The subordinate distributions characterized the fine-scale structures within each category. Samples drawn from each subordinate distribution corresponded to brain word representations (for more details, see Methods). We first defined a baseline parameter set for the Gaussian distributions that simulates the structures of putative healthy controls (Fig. 5a). Then, categorical and within-category structures were manipulated by changing these parameters from baseline to yield structures of possible cases of patients. We varied the following three parameters of these Gaussian distributions separately: the variance of mean of the superordinate distributions (case 1), the variance of mean of the subordinate distributions (case 2), and the variance of samples from the subordinate distributions (case 3 in Fig. 5b–d). In case one, categorical structure is changed whereas in cases 2 and 3 within-category structure is changed.

We found that only a change in the variance of samples from the subordinate distributions (case 3) was needed to replicate the fMRI data (Fig. 5e and f; see also Supplementary Fig. 9). When the sample variance of each subordinate distribution was increased (Fig. 5d) compared to putative controls (Fig. 5a), the variance in word-pairwise distances of both overall and within-category brain word representations decreased. Our additional simulation also suggests that categorical structure without any fine-scale structure in brain word representations is insufficient to explain the observed fMRI results (Supplementary Fig. 10). These results raise the possibility that disrupted within-category fine-scale structures of semantic representations underlie the reduced representational variability in schizophrenia patients.

### Associations of variability reductions with psychopathological symptoms

To examine if this reduced representational variability is associated with pathological symptoms in schizophrenia, we calculated Spearman’s correlations of these variance measures with scores on the 5-factor Positive and Negative Symptom Scale^25^ (PANSS-5, “positive”, “negative”, “disorganized”, “excitement”, “depressive”, and “total”). The two intra-individual variability measures word-pairwise distance and reduced-dimensional representation consistently showed strongest correlations with the “disorganized” factor (Fig. 5a and b), although the correlations fell below statistical significance after Bonferroni correction for multiple comparisons. According to the original study on the PANSS-5, the “disorganized” factor is closely linked with conceptual disorganization in schizophrenia^25^. Therefore, these results support the notion that the observed reductions of intra-individual variability in brain word representations reflect the disorganization of semantic representations.

**Fig. 6.**
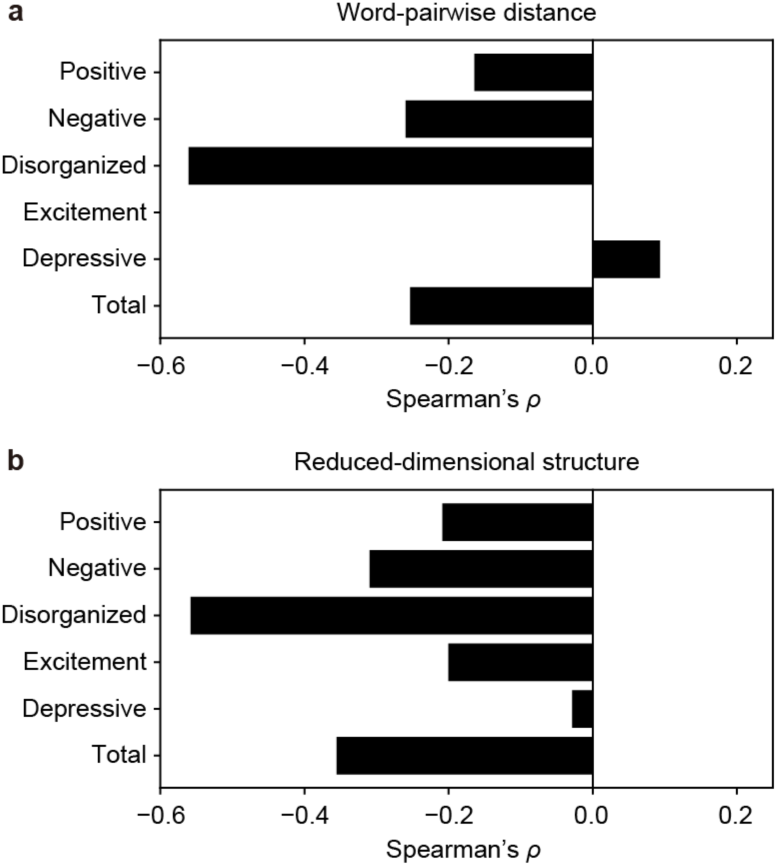
Relationship between representational variability reduction and psychopathological syndromes of schizophrenia. **a**, The correlation of intra-individual variability in word-pairwise distances with PANSS-5 factors. Spearman’s correlation coefficients (*ρ* values) between the variance of word-pairwise distances and each of the 5 PANSS factors were calculated across patients and revealed strongest associations with the PANSS factor “disorganization”. **b**, The correlation of intra-individual variability in reduced-dimensional representations with PANSS-5 factors. The explained variance ratio of the 1st PCs of brain word representations was also most strongly associated with “disorganization”.

## Discussion

We quantified the structural properties of semantic representations in individual schizophrenia patients using voxelwise modeling with a word vector space and found lower variability both within and across individuals compared to healthy controls. This reduced representational variability in patients was observed even in the categorical structures of semantic representations. To our knowledge, this is the first study demonstrating disorganization of semantic information in schizophrenia by directly evaluating brain semantic representations. In addition, reduced representational variability was associated with a psychopathological symptom scale reflecting conceptual disorganization. Our findings suggest that reductions in intra- and inter-individual representational variability are associated with the semantic disorganization characteristic of schizophrenia.

Intra-individual variability was reduced not only across all representations (Fig. 2) but also within semantic categories (Fig. 4a), and the observed variability reduction was replicated by simulating the disruption of semantic representations within individual categories (Fig. 5). Such disruption could impair the ability to use semantic associations between words to guide cognition and behavior. In fact, such a deficit may explain the abnormalities observed in a wide range of behavioral tests, such as verbal fluency^2,26^ and semantic priming^27,28^. Several previous behavioral studies have even suggested that disorganization of semantic storage in the brain is a key contributor to semantic processing impairments^8–11^. The present study provides neurophysiological evidence to support this hypothesis and provides clues to the nature of semantic storage disorganization at the level of brain representations.

Previous studies have also reported reduced intra-individual variability in the dynamic properties of resting-state brain activity among schizophrenia patients^29,30^. For example, Miller et al. found that resting-state connectivity patterns are markedly less variable over time among schizophrenia patients compared to healthy controls^29^. Therefore, reduced intra-individual variability may be a central pathogenic characteristic of both resting-state and stimulus-evoked neural processing in schizophrenia.

The present study also demonstrated reduced inter-individual variability in semantic representations across schizophrenia patients (Figs. 3, 4c and 4d). This reduction may be also induced by the disruption of within-category semantic structure. We speculate that this disruption diminishes the uniqueness of within-category semantic structures that individuals intrinsically have, leading to reduced inter-individual semantic variability. Such reduced variability may produce cognitive and behavioral abnormalities typical of schizophrenia.

Reduced inter-individual variability among schizophrenia patients was also reported in a large-scale investigation of resting-state fMRI data^31^. By quantifying sampling bias (differences between subject populations) in individual resting-state functional connectivity independently of other bias factors, they showed that the sampling bias is smaller among patients than healthy controls. Therefore, like reduced intra-individual variability, reduced inter-individual variability may be a key characteristic of both resting-state and stimulus-dependent neural processing in schizophrenia.

One possible reason for reduced inter-individual representational variability is shared genetic variants among patients with schizophrenia, which is known to be highly heritable^32^. Consistent with this notion, previous studies have identified several genetic variants associated with schizophrenia^33–37^, some of which are also related to brain structure^38–40^. In turn, brain structure is related to semantic processing^41–43^. If the genetic factors underlying schizophrenia are directly or indirectly associated with the organization of semantic representations in the brain, it is possible that similar maladaptive organizational patterns are shared among schizophrenia patients harboring common genetic anomalies. It will be of interest to investigate the shared genetic basis of schizophrenia and semantic organization.

This reduced variability in semantic representations may be useful for the clinical diagnosis of schizophrenia. Conventionally, semantic impairments in schizophrenia are diagnosed using tests of synonym generation^8,44^, picture naming^7,44^, and verbal fluency^9,45^. However, these methods do not directly evaluate the structural properties of semantic representations, although verbal fluency can be used to indirectly estimate the semantic organization of a small number of word representations^10,11,14^. If semantic disorganization is a shared feature among schizophrenic, direct evaluation of structural properties may be a more efficient tool to diagnose semantic impairments. The pairwise distances of object representations in the brain can be evaluated using a sophisticated psychological test ^46,47^, and we suggest that this same test could be used to estimate the pairwise distances of word representations in the schizophrenic brain. Based on test results, we may be able to diagnose schizophrenia by assessing word-pairwise distances and compared intra-individual representational variability to a normal standard. Further investigations will be needed to provide a proof-of-concept for the diagnostic utility of this method.

## Methods

### Subjects

Fourteen patients with schizophrenia (age 24–59 years, mean = 44.1 years; 6 females) participated in this study. Each patient fulfilled the criteria for schizophrenia based on the Structured Clinical Interview for DSM-IV Axis I Disorders (SCID) Patients Edition version 2.0. All patients were free of comorbid mental disorders. Patients were also examined using the Japanese Version of the National Adult Reading Test (JART) short form, which is thought to reflect premorbid IQ^48^. Psychopathology was assessed using the 5-factor Positive and Negative Symptom Scale^25^ (PANSS-5) and Peters et al Delusions Inventory (PDI)^49^. Seventeen adults (age 26–49 years, mean = 40.2 years; 10 females) with no psychopathology as evaluated using the SCID Non-patient Edition version 2.0, no history of psychiatric disorders, and no psychosis in first-degree relatives were recruited as controls. They were matched to patients for age, gender, handedness, and predicted IQ.

All subjects had normal or corrected-to-normal vision. They provided written informed consent after receiving a complete description of the study. The experimental protocol was approved by the Committee on Medical Ethics of Kyoto University and conducted in accordance with The Code of Ethics of the World Medical Association.

### MRI data collection

Functional and anatomical MRI data were collected using a 3T Siemens TIM Trio scanner (Siemens, Germany) and a 32-channel Siemens volume coil. Functional data were collected using a gradient echo EPI sequence (TR = 2000 ms; TE = 30 ms; flip angle = 70°; voxel size = 2.5 × 2.5 × 3.6 mm; matrix size = 80 × 80; the number of slices = 37). Anatomical data were collected using a T1-weighted MPRAGE sequence (TR = 2000 ms; TE = 3.4 ms; flip angle = 8°; voxel size = 0.94 × 0.94 × 1.0 mm; matrix size = 240 × 256; the number of slices = 208) on the same 3T scanner.

### Experimental design

The visual stimuli consisted of silent color movie clips (10–20 s each) of natural scenes taken from movies used in previous studies^18,50^ and spliced together in random order. The subjects viewed the clips on a projection screen in the MRI scanner while they freely moved their gaze. Subjects were scheduled to view six 610-s spliced movies in separate scans, but 3 of the 14 patients viewed only 4. In each movie, the initial 10-s was a dummy to discard hemodynamic transients caused by movie onset and not used for analysis. The following 60-s was identical across all six movies, while the remaining clips were different. Responses during these latter clips were used for estimating the parameters of voxelwise models (training dataset; see voxelwise modeling), while responses to the former clip were used to validate the fitted models (test dataset).

### fMRI data preprocessing

Motion correction in each functional scan was performed using the statistical parameter mapping toolbox (SPM8, http://www.fil.ion.ucl.ac.uk/spm/software/spm8/). For each subject, all volumes were aligned to the first image from the first functional run. Low-frequency fMRI response drift was eliminated by subtracting median-filtered signals (with a 120-s window) from raw signals. Then, the response for each voxel was normalized by subtracting the mean response and scaling to the unit variance. FreeSurfer^51,52^ with the Destrieux atlas^53^ was used to identify cortical surfaces from anatomical data and register these surfaces to the voxels of functional data. All voxels identified within the whole cortex and subcortical structures (dorsal and ventral striatum, pallidum, amygdala, hippocampus, thalamus, brain stem, and cerebellum) of each subject were used for voxelwise modeling.

### Semantic vector construction for movie scenes

Scene descriptions were given for every 1-s of the movie by native Japanese speakers who were not the fMRI subjects. These annotators were instructed to describe each scene using more than 50 Japanese characters (see Fig. 1a for an example). Five annotators were randomly assigned for each scene to reduce the potential effects of personal bias. These descriptions were then transformed into vectors using word2vec^22^ as described previously^17^. The word2vec (skip-gram) algorithm was originally developed to learn a high-dimensional word vector space based on local (nearby) word co-occurrence statistics in natural language texts. The vector space effectively captures the semantic relationships among many words and can be used to model the semantic representations in the human brain^16,21^. In the present study, a word2vec vector space was preconstructed from a text corpus of the Japanese Wikipedia dump on January 11, 2016 (vector dimensionality = 1000; window size = 10; the number of negative samples = 5). All Japanese texts in the corpus were segmented into words and lemmatized using MeCab (http://taku910.github.io/mecab). Only nouns, verbs, and adjectives were used. To improve the reliability of word2vec learning and restrict the vocabulary size to around 100000 words, words that appeared less than 178 times in the corpus were excluded.

Each description for a given scene was also segmented, lemmatized, and decomposed into nouns, verbs, and adjectives using MeCab as described above and transformed to word2vec vectors. The word vectors were then averaged within each description. For each scene, all vectors obtained from the different descriptions were averaged. Through this procedure, one 1000-dimensional vector (scene vector) was obtained for each 1-s scene and used for the modeling.

### Voxelwise modeling

A series of the response **r** evoked in each of *N* voxels by a series of *S* movie scenes was modeled as a weighted linear combination of a scene-vector matrix **V** plus isotropic Gaussian noise **ε**.

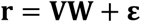

A set of linear temporal filters was used to model the slow hemodynamic response and its coupling with brain activity^50^. To capture the hemodynamic delay in the responses, the *S* × 3*K* matrix **V** was constructed by concatenating three sets of *K*-dimensional scene vectors with delays of 2, 4, and 6 s. The 3*K* × *N* weighted matrix **W** was estimated using an L2-regularized linear least-squares regression, which can obtain good estimates even for models containing a large number of regressors^18,54^.

To estimate the regularization parameters, the training dataset was randomly divided into two subsets containing 80% and 20% of samples for model fitting and validation, respectively. This random resampling procedure was repeated 10 times. Regularization parameters were optimized according to the mean Pearson’s correlation coefficient between the predicted and measured BOLD signals for the 20% validation samples. The optimal parameter was obtained separately for each subject and for each dimension of scene vectors.

The brain-response prediction performance of the model was evaluated using the test dataset, which was not used for model fitting or parameter estimation. The prediction accuracy of the model was quantified as the Pearson’s correlation coefficient between the predicted and average measured BOLD signals in the test data set.

### Brain word representation

The representations of individual words in each subject’s brain were quantified using the weights of that subject’s voxelwise model. First, the 3*K* × *N* weight matrix was transformed into *K* × *N* by averaging the weights across the three sets of hemodynamic delay terms. Next, the top *M* (= 7000) voxels with highest prediction accuracy were selected for making the *K* × *M* weight matrix. Finally, the original *K*-dimensional word2vec vectors of the top 2000 most frequently appearing words in the movie scene descriptions were multiplied by the *K* × *M* weight matrix, yielding *M*-dimensional vectors of 2000 words. These vectors were obtained for each subject and regarded as brain word representations of the subject.

### Structural properties of brain word representations

The structural properties of brain word representations within and across subjects were evaluated using word-pairwise distance and reduced-dimensional representation. The word-pairwise distances for a given subject reflect the structure of semantic dissimilarity between each pair of brain word representations in that subject. They were evaluated by the correlation distances between all possible pairs of brain word representations for that subject. The correlation distance was computed by 1 – *r*, where *r* is a Pearson’s correlation coefficient, and thereby ranged from 0 to 2. The variance of all word-pairwise distances for each subject was used to evaluate the intra-individual variability of semantic representations. Then, the subject-pairwise correlation of word-pairwise distances was computed by the Pearson’s correlation between all word-pairwise distances for one subject and all corresponding word-pairwise distances for other subjects within the group. The subject-pairwise correlations for all possible subject pairs within each group were used to evaluate the inter-individual variability of structural properties of brain word representations.

The reduced-dimensional representations were estimated using principle component analysis (PCA) of brain word representations. The brain word representations of 2000 words for each subject were represented by 7000 voxels. The number of words was regarded as dimensions and reduced using PCA. This produced 2000-dimensional vectors as principle components (PCs). These vectors reflect how semantic representations are distributed in particular dimensions after dimensional reduction. Since a higher explained variance ratio of a given PC indicates higher variance in brain word representations along that PC, the intra-individual variability of semantic representations was evaluated by the explained variance ratio of each PC. Then, the subject-pairwise correlations of the reduced-dimensional representations were evaluated by the Pearson’s correlations between each PC for one subject and the corresponding PC of all other subjects within the group. Since PCA assumes that two PCs with opposite directions (180°) are identical, negative values of the subject-pairwise correlations (>90°) were transformed into positive values (≤ 90°); that is, the absolute values were used. The absolute values of subject-pairwise correlations of all possible subject pairs within each group were used to evaluate the inter-individual variability of structural properties.

### Dimensional shuffling of brain word representations

To examine if brain word representations are intrinsically structured, dimensional shuffling was performed with values along the dimensions of *M* (=7000; i.e., the number of voxels). In dimensional shuffling, the values were randomly shuffled across all representations separately for each dimension. This manipulation produced a new set of brain word representations with altered structure but preserved variance in each dimension. Then, the structural property measures word-pairwise distance and reduced-dimensional representation were re-examined. This procedure was repeated 100 times. Finally, a set of 100 structural property measures were averaged separately for the two types of measures. The decrease in structure after dimensional shuffling was regarded as the degree of intrinsic structure in the brain word representations.

### Manual word categorization

To examine the categorical structures of semantic representations, six categories of words were manually created, “human”, “(non-human) animal”, “(non-animal) natural thing”, “body part”, “building”, and “vehicle”, each of 25 unique words (see also Supplementary Table 2). According to previous reports, semantic representations related to humans, animals, and vehicles form separate clusters in the brain representational space^18^. In addition, the human object-selective cortex shows clear selectivity for information on body parts and buildings^55,56^. Hence, it would be expected that brain word representations of these categories normally form cluster-like structures in the representational space.

### Automatic word categorization using unsupervised clustering

The initial clustering analysis used words arbitrarily chosen for each semantic category. However, this process cannot completely exclude bias. To eliminate potential bias, words were also categorized using an unsupervised clustering method, x-means^24^, which is an extension of k-means. The x-means clustering automatically estimates not only the cluster structure of a given dataset by k-means but also the optimal number of clusters for the dataset on the basis of the Bayesian information criterion^57^. The x-means clustering with k-means++ initialization^58^ was applied to 2000 brain word representations for each subject to estimate own word categories. Note that since the word assignment of this categorization varied across individuals, the inter-individual similarity of categorical representations could not be tested.

### Computer simulations of representational variability reduction

Computer simulations were performed to provide a possible explanation for the group differences in representational variability (Figs. 2 and 4) observed in the fMRI dataset. In the simulation, brain word representations were assumed to have both categorical structures (i.e., separation into categories) and fine-scale structure within each category as shown by dimensional shuffling analyses on fMRI data (Supplementary Fig. 6). This two-level structure was mimicked by nested Gaussian distributions separated into (1) superordinate distributions that characterized the categorical structure and (2) subordinate distributions that characterized the within-category fine-scale structure (Fig. 5a). Three subordinate distributions were included in each of the six superordinate distributions and had different mean parameters drawn from the corresponding superordinate distribution. Samples drawn from each of the subordinate distributions corresponded to the brain word representations.

Each Gaussian distribution was represented by the formula

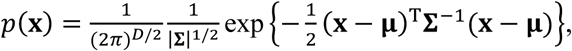

where *D* denotes the dimensionality of a representational space, **μ** denotes the mean, and **Σ** denotes a covariance matrix. For computational efficacy, D = 100 was used, although the original dimensionality of brain word representations was 7000. For simplicity, the covariance matrix was a scalar matrix represented by **Σ** = *a***I**, where *a* is a scalar value and **I** is a unit matrix.

The procedure for this simulation was as follows: (1) The mean values of six superordinate distributions (**μ**_1_) were randomly drawn from a Gaussian distribution with mean of 0 and variance of *a*_0_**I**; (2) The mean values of the three subordinate distributions (**μ**_2_) were randomly drawn from the superordinate distributions with different mean (**μ**_1_ value) but shared covariance matrix (*a*_1_**I**); (3) Brain word representations (50 samples for each subordinate distribution) were randomly drawn from each of the subordinate distributions with different **μ**_2_ values and a shared covariance matrix (*a*_2_**I**); (4) Representational variability measures (the variability of word-pairwise distance) were computed from the simulated brain word representations. This procedure was repeated 1000 times. Finally, the set of 1000 measures was averaged. The three parameters *a*_0_, *a*_1_, and *a*_2_ determine the structural variability of brain word representations. As a baseline parameter set (corresponding to the structure for healthy controls), an *a*_0_ of 1.0, *a*_1_ of 1.4, and *a*_2_ of 1.0 were used. The structural changes of brain word representations in putative patients were simulated by varying each of the three parameters from 0.2 to 2.6 in steps of 0.4 while keeping the other parameters fixed (Supplementary Fig. 9). The representative simulation results (structural changes due to variation of the three parameters) were obtained using *a*_0_ of 0.6, *a*_1_ of 1.4, and *a*_2_ of 1.0 for case 1, *a*_0_ of 1.0, *a*_1_ of 1.0, and *a*_2_ of 1.0 for case 2, and *a*_0_ of 1.0, *a*_1_ of 1.4, and *a*_2_ of 2.2 for case 3.

## Acknowledgments

The work was supported by JSPS KAKENHI Grant-in-Aid for Early-Career Scientists (18K18141 and 19K17061), for Young Scientists A (15H05311), for Scientific Research B (17H01797, 19H03583, and 19H04200), for challenging Exploratory Research (16K13106), and 16K13117), and Research on Innovative Areas (16H06572, 18H04208, and 18H05019), JST CREST (JPMJCR18A5), and JST ERATO (JPMJER1801) from MEXT and by a grant from Tateishi Science and Technology Promotion Foundation (2191025). This work was based on results obtained commissioned by NEDO. A part of this study is the result of the Strategic Research Program for Brain Sciences (19dm0307008h0002, 19dm0107151s0104) by the Japan Agency for Medical Research and Development (AMED) and research and development of technology for enhancing functional recovery of elderly and disabled people based on non-invasive brain imaging and robotic assistive devices, the Commissioned Research of National Institute of Information and Communications Technology, JAPAN.

## Author contributions

S. Nishida, Y.M., S. Nishimoto, and H.T. designed the research. Y.M., S.S., and A.M. performed experiments. S. Nishida and Y.N. performed the analysis. S. Nishida, Y.M., N.Y., S.S., A.M., R.H., S. Nishimoto, and H.T. wrote the manuscript.

---

**Supplementary Fig. 1.**
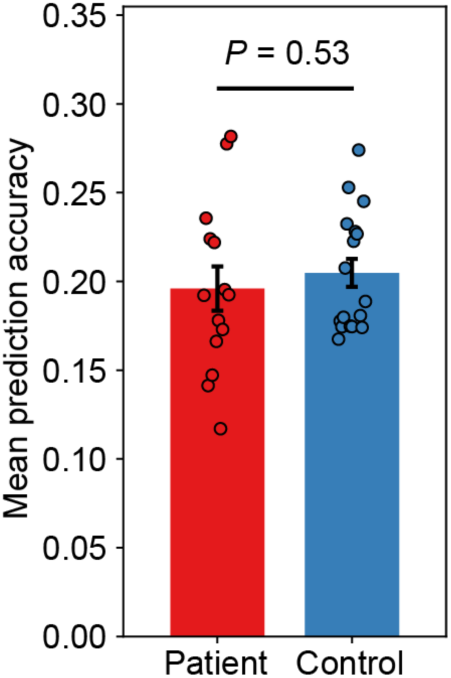
Comparison of response-prediction accuracy between subject groups. For each subject, response-prediction accuracy of the voxelwise model was evaluated by the Pearson’s correlation coefficient between predicted and measured voxel response. The evaluation was performed with the test dataset, which was not used for model training. The 7,000 voxels with greatest accuracy (highest correlation) were used for calculating brain word representations. The mean accuracy did not differ between patients (red) and controls (blue) (Mann-Whitney U test, P = 0.53). Bars depict the mean values for each subject group (error bars, SEM). Each circle depicts a single subject.

**Supplementary Fig. 2.**
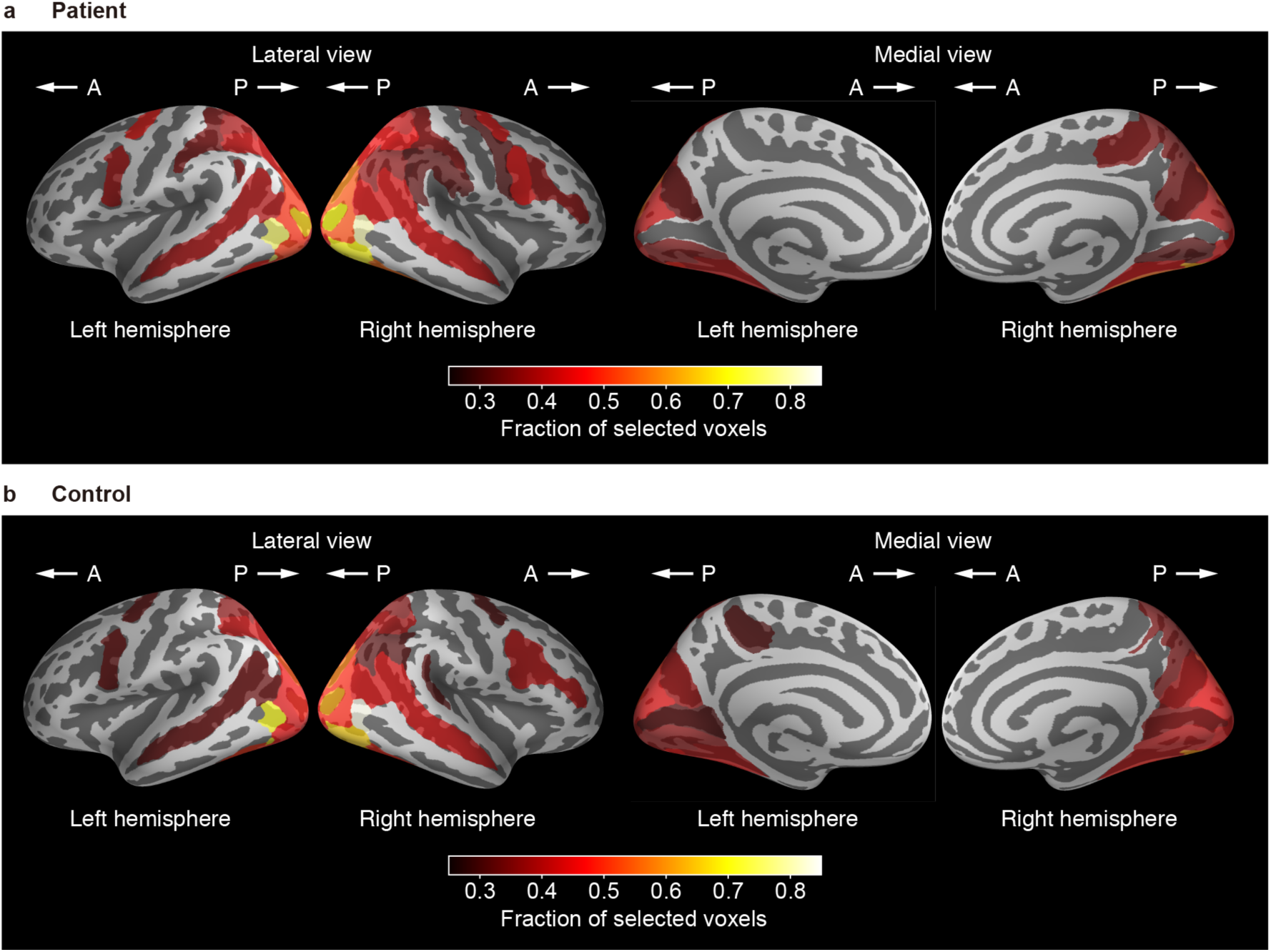
Cortical surface maps showing the fraction of voxels selected for calculating brain word representations in each anatomical region-of-interest (ROI). To compare the localization of voxels selected for calculating brain word representations (i.e., 7,000 voxels with the highest response-prediction accuracy) between groups, the fraction of selected voxels was mapped onto the segmented cortical surface of a reference brain. For this purpose, the cortex of each subject was anatomically segmented into 150 ROIs using FreeSurfer with the Destrieux atlas. For each ROI, the fraction of selected voxels to all voxels in the ROI was computed and averaged across patients (**a**) or controls (**b**). Brighter locations in the surface maps indicate ROIs with higher fractions. Only ROIs with fractions above 0.25 are shown. Abbreviations in the figure: A, anterior; P, posterior. See also Supplementary Table 1.

**Supplementary Fig. 3.**
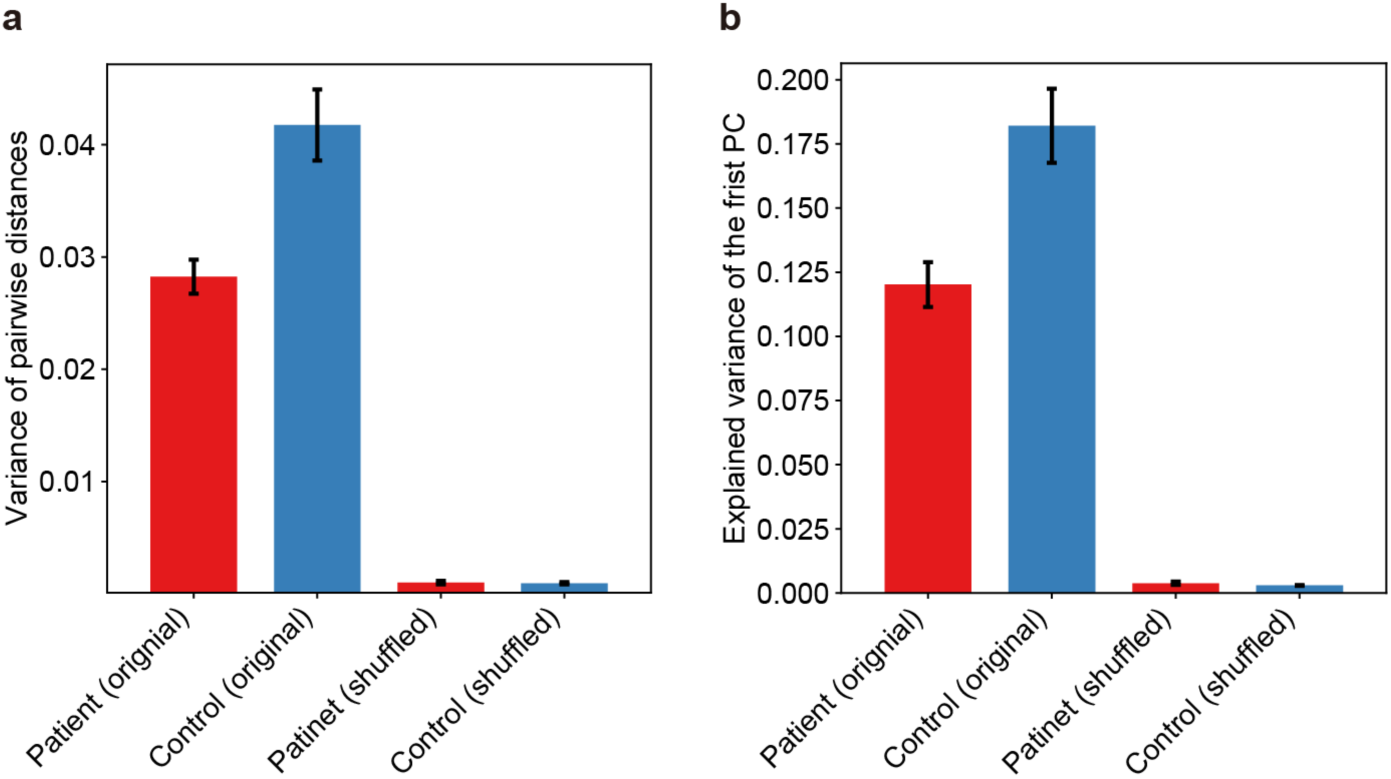
Intra-individual representational variability after dimensional shuffling. To test whether brain word representations are randomly distributed or intrinsically structured, we compared structural properties after the values of brain word representations were randomly shuffled across all representations for each dimension. The variance of pairwise distances (**a**) and the explained variance of the first PC (**b**) were computed using original (two left bars) and dimensionally shuffled (two right bars) brain word representations in patients (red) and controls (blue). Bars depict the mean values for each subject group (error bars, SEM). After dimensional shuffling, the structural properties were reduced in both groups (Wilcoxon test, p < 0.001), suggesting that brain word representations are not randomly distributed but rather structured in both groups.

**Supplementary Fig. 4.**
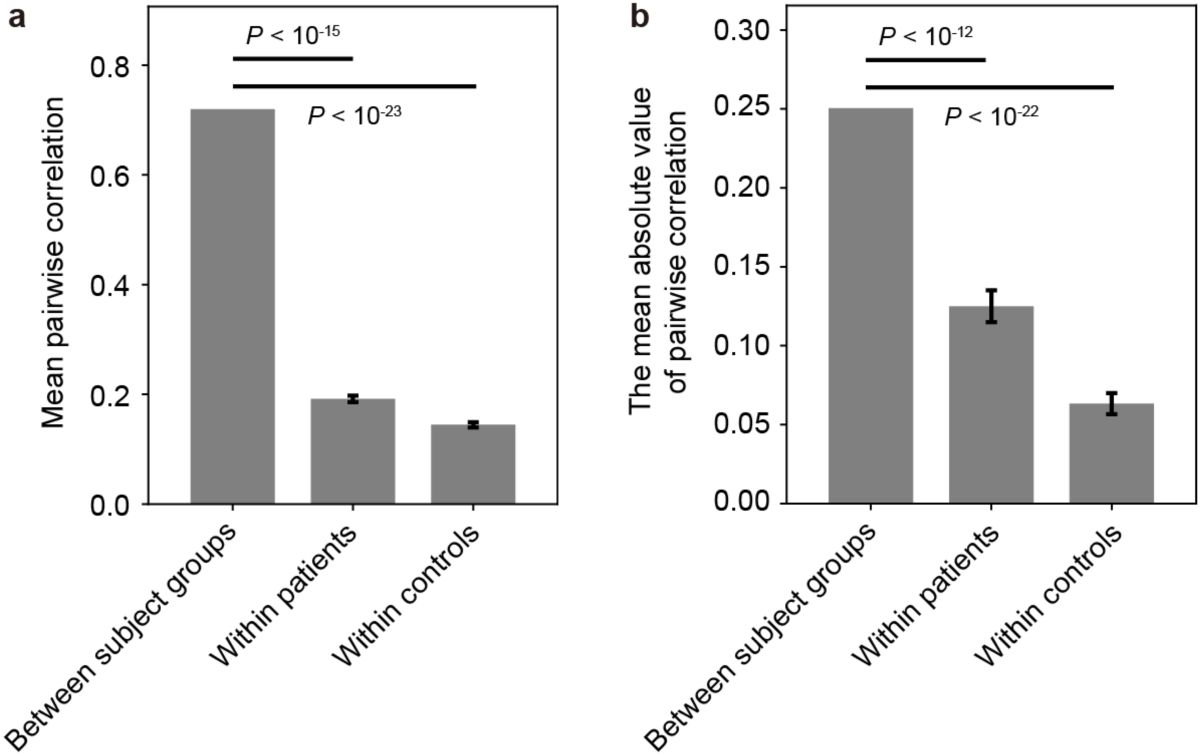
Correlations of structural properties within and between groups. To test whether there are distinctive structural differences in brain word representations within and between groups, we calculated the correlations of mean structural properties between all patient–patient, control–control, and patient–control subject pairs (Fig. 3). For the word-pairwise distances (**a**), the mean correlation between controls and patients was significantly stronger than that between patients (Wilcoxon test, P < 10^−15^) and between controls (P < 10^−23^). Error bars are SEM. For the reduced-dimensional representations (**b**), the mean correlation between controls and patients was significantly stronger than that between patients (P < 10^−12^) and between controls (P < 10^−22^). Hence, the inter-individual variabilities of these two structural property measures were much lower between groups than within groups. This indicates that patients do not have structure properties clearly distinct from controls, at least at the group-average level.

**Supplementary Fig. 5.**
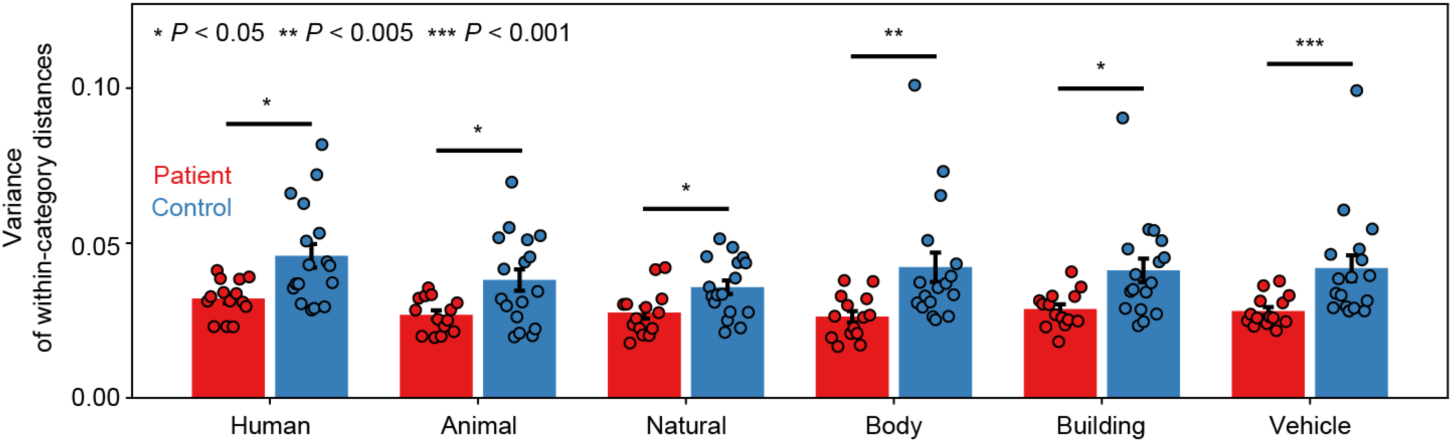
Category-specific comparisons of intra-individual variability. For each of the six word categories, the within-subject variance was computed and compared between subject groups. Same conventions as Fig. 4**a**. The within-subject variance significantly differed for all categories (Mann-Whitney U test, P < 0.05). Marks above each group of bars denote the significance of the subject-group comparisons (*P < 0.05; **P < 0.005; ***P < 0.001).

**Supplementary Fig. 6.**
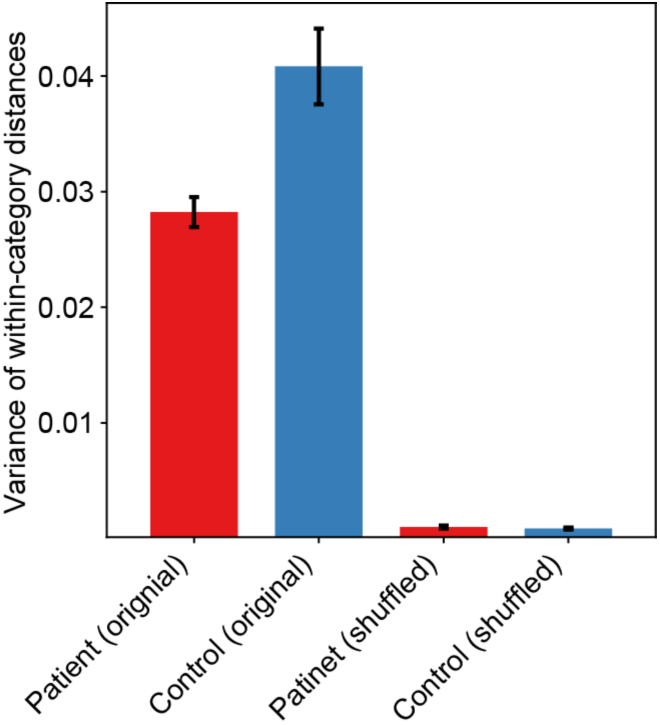
Intra-individual variability of within-category representations after dimensional shuffling. We compared the within-subject variance of within-category word pairwise distances at baseline (original) and after dimensional shuffling of brain word representations. The shuffling procedure was the same as used for overall brain word representations (Supplementary Fig. 3) except that shuffling was limited to brain word representations contained in individual categories. Same conventions as Supplementary Fig. 3. The variance drastically decreased for both patients and controls after dimensional shuffling (Wilcoxon test, p < 0.001), suggesting that brain word representations within semantic categories are not randomly distributed in either group.

**Supplementary Fig. 7.**
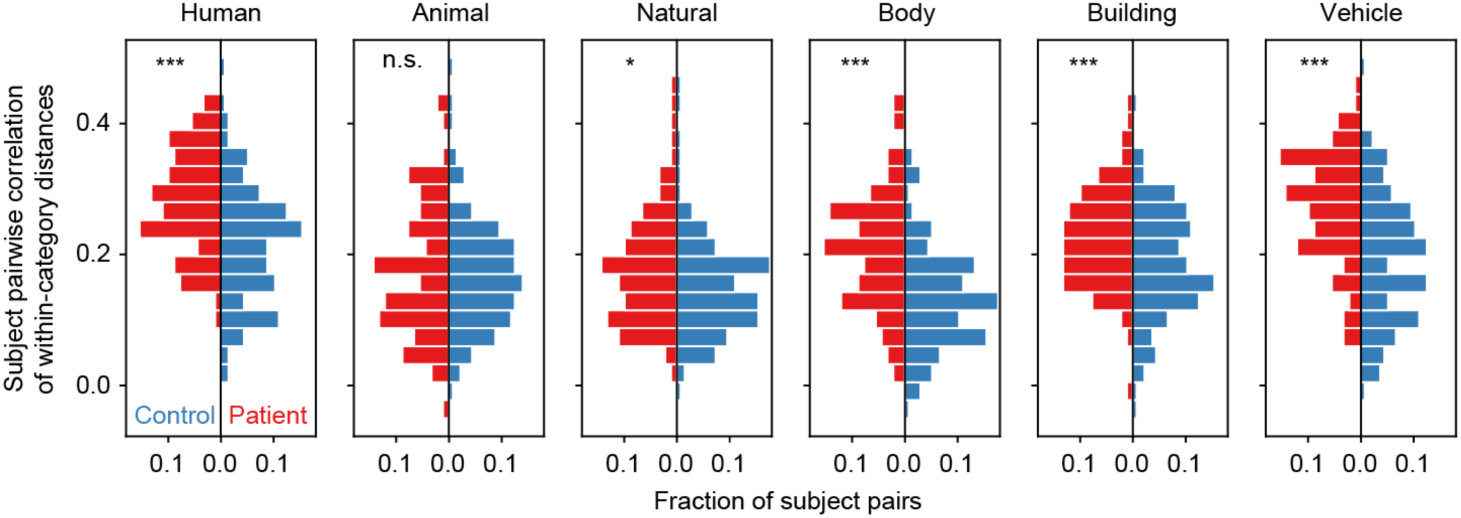
Category-specific comparisons of inter-individual variability in word-pairwise distances. For each of the six word categories, the subject-pairwise correlations were computed and compared between subject groups. Same conventions as Fig. 4**c**. The subject-pairwise correlations differed significantly for all semantic categories except for “animal” (P < 0.05). Marks in the top left of each panel denote the significance of the subject-group comparisons (*P < 0.05; **P < 0.005; ***P < 0.001; n.s, P > 0.05).

**Supplementary Fig. 8.**
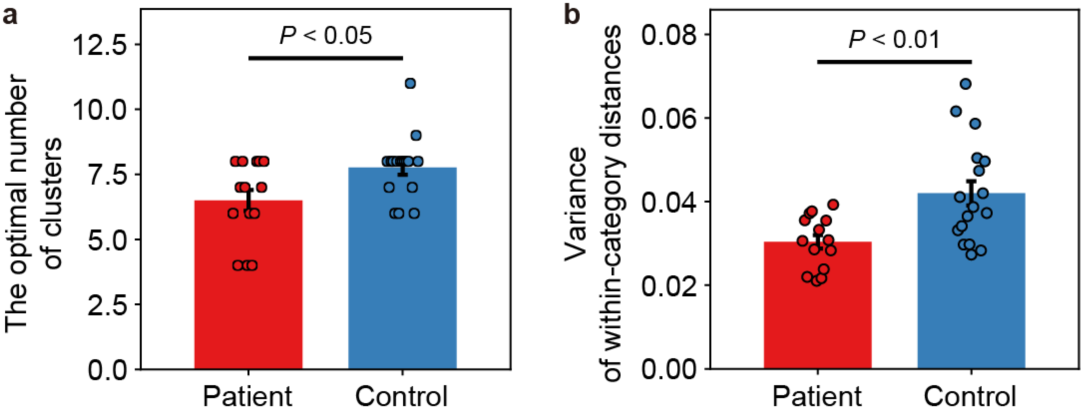
Representational variability within categories estimated by unsupervised clustering. To eliminate the possibility that the observed reduced representational variability within individual categories is caused by arbitrary word selection for manual categorization, we also categorized brain word representations by x-means clustering, which is an extension of k-means clustering with automatic estimation of the optimal number of clusters for target datasets. **a**, The optimal number of clusters estimated by x-means was significantly smaller for patients than for controls (Mann-Whitney U test, P < 0.05). **b**, The variance of within-category distances between brain word representations, evaluated from the pairwise distances between all possible pairs within each estimated cluster, was significantly smaller among patients than controls (P < 0.01). Same conventions as Fig. 4**a**.

**Supplementary Fig. 9.**
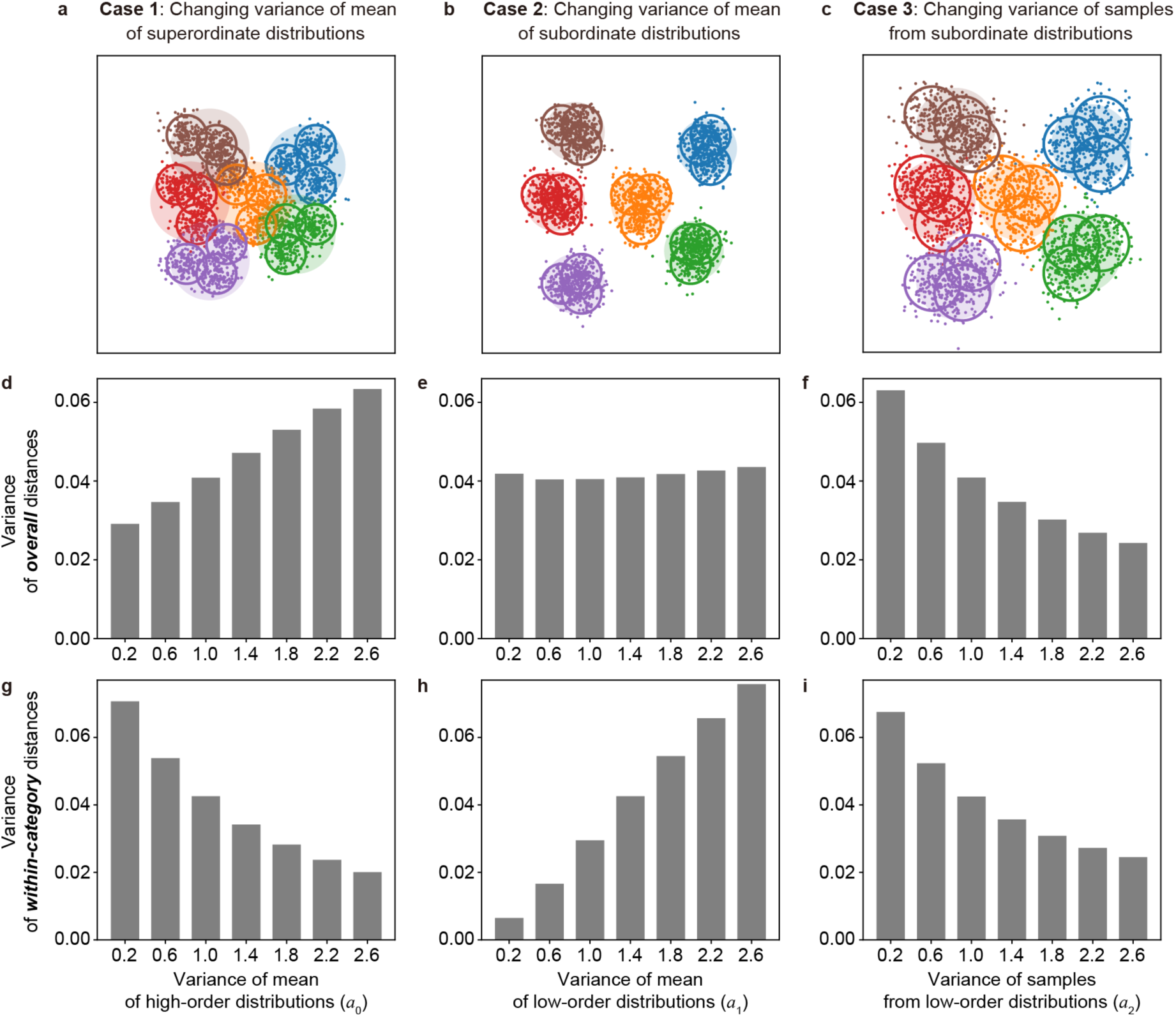
Effects of three variance parameters on representational variability in computer simulations. **a**–**c**, Computer simulations of the three parametric changes (cases 1–3) shown in Fig. 5**b**–**d**, changes in variance of the mean of the superordinate distributions (case 1), changes in the variance of the mean of subordinate distributions (case 2), and changes in the variance of samples from subordinate distributions (case 3). We examined the changes in the variance of overall word-pairwise distances (**d**–**f**) and within-category word-pairwise distances (**g**–**i**) of brain word representations. In case 1 (**d** and **g**; *a*_0_=0.2–2.6, *a*_1_=1.4, *a*_2_=1.0), the two variance types changed in opposite directions with the change in *a*_0_. In case 2 (**e** and **h**; *a*_0_=1.0, *a*_1_=0.2–2.6, *a*_2_=1.0), the variance of overall distances was relatively unchanged under changing *a*_1_. In case 3 (**f** and **i**; *a*_0_=1.0, *a*_1_=1.4, *a*_2_=0.2–2.6), the two variances changed in the same direction with changing *a*_2_. Therefore, only in case 3 could we obtain a variance reduction pattern consistent with our fMRI data (see also Fig. 5).

**Supplementary Fig. 10.**
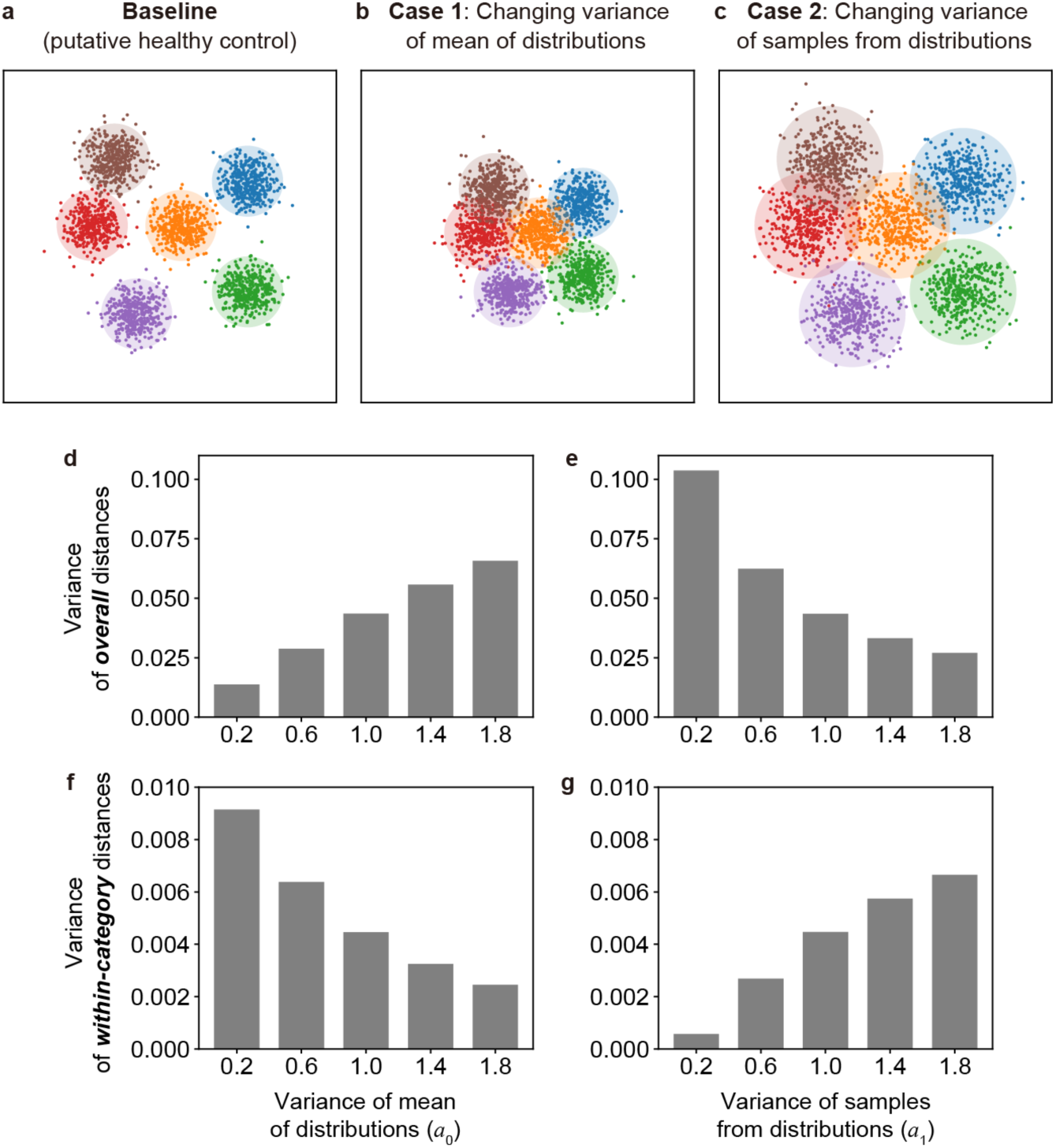
Effects of variance parameters on representational variability in computer simulations with no within-category structures. **a**–**c**, Computer simulation based on the assumption that brain word representations are not organized with fine-scale structure within categories. In this simulation, samples corresponding to brain word representations were generated directly from six Gaussian distributions in a 100-dimensional space and did not use the nested Gaussian distributions (**a**). The mean parameters ***μ***_1_ of the six distributions were randomly drawn from a Gaussian distribution with the mean of 0 and variance of *a*_0_**I**. Then, brain word representations (150 samples for each distribution) were randomly drawn from each of the distributions with different ***μ***_1_ values and a shared covariance matrix (*a*_1_**I**). We manipulated structural changes by varying the two parameters *a*_0_ (**b**; case 1) and *a*_1_ (**c**; case 2) separately and examined their effects on variance of overall word-pairwise distances (**d** and **e**) and within-category word-pairwise distances (**f** and **g**) of brain word representations. In case 1 (**d** and **f**; *a*_0_ = 0.2–1.8, *a*_1_ = 1.0) and case 2 (**e** and **g**; *a*_0_ = 1.0, *a*_1_ = 0.2–1.8), the two variance types changed in opposite directions as each of the parameters was changed. This indicates that categorical structure changes in a brain word representation matrix without within-category (fine-scale) structure cannot replicate the pattern of intra-individual variability reduction observed by fMRI.

**Supplementary Table 1.**
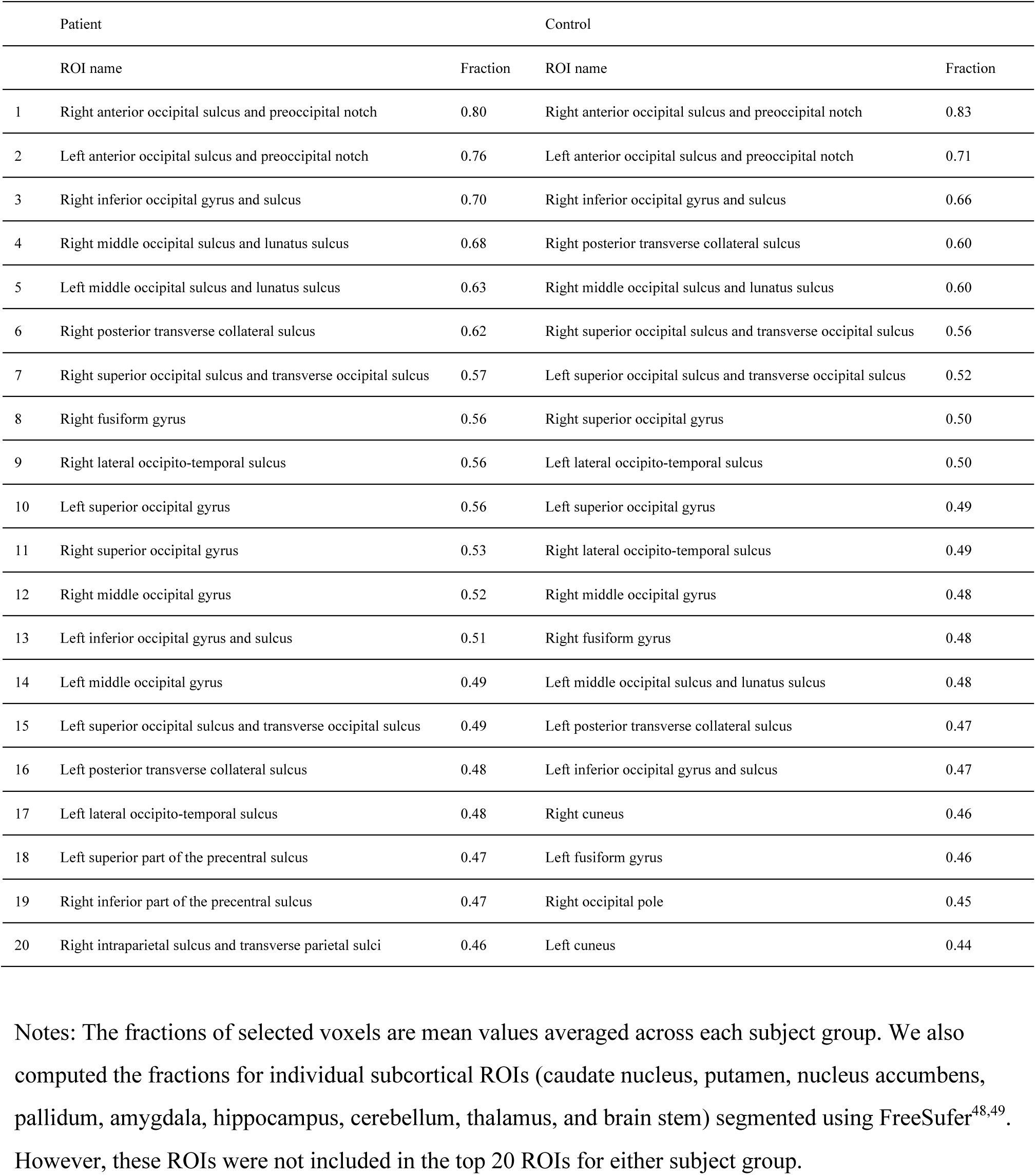
The 20 anatomical regions of interest (ROIs) with the greatest fractions of voxels selected for calculating brain word representations.

**Supplementary Table 2.**
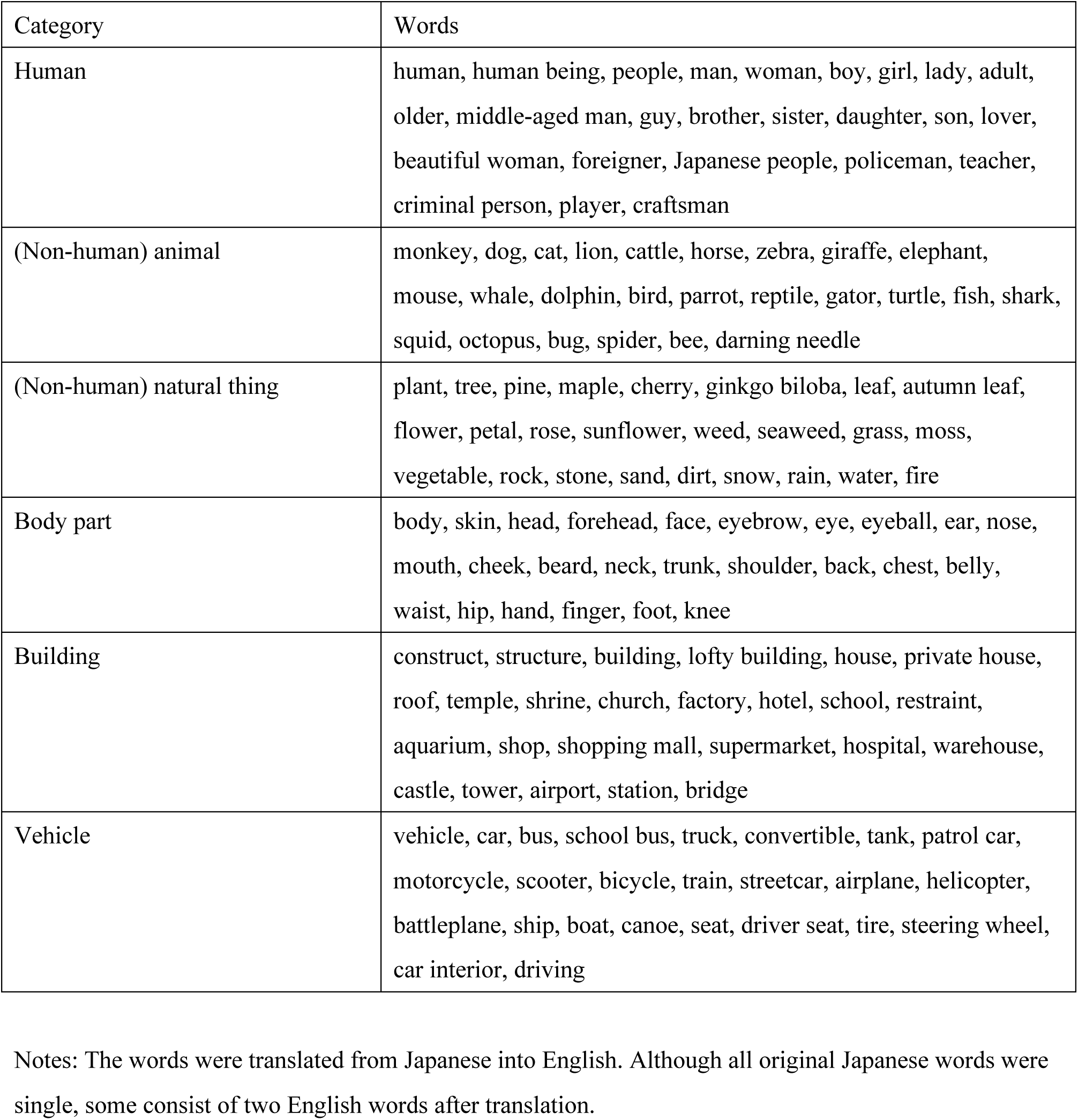
25 words belonging to each of the 6 word categories.

| Category | Words |
| --- | --- |
| Human | human, human being, people, man, woman, boy, girl, lady, adult, older, middle-aged man, guy, brother, sister, daughter, son, lover, beautiful woman, foreigner, Japanese people, policeman, teacher, criminal person, player, craftsman |
| (Non-human) animal | monkey, dog, cat, lion, cattle, horse, zebra, giraffe, elephant, mouse, whale, dolphin, bird, parrot, reptile, gator, turtle, fish, shark, squid, octopus, bug, spider, bee, darning needle |
| (Non-human) natural thing | plant, tree, pine, maple, cherry, ginkgo biloba, leaf, autumn leaf, flower, petal, rose, sunflower, weed, seaweed, grass, moss, vegetable, rock, stone, sand, dirt, snow, rain, water, fire |
| Body part | body, skin, head, forehead, face, eyebrow, eye, eyeball, ear, nose, mouth, cheek, beard, neck, trunk, shoulder, back, chest, belly, waist, hip, hand, finger, foot, knee |
| Building | construct, structure, building, lofty building, house, private house, roof, temple, shrine, church, factory, hotel, school, restraint, aquarium, shop, shopping mall, supermarket, hospital, warehouse, castle, tower, airport, station, bridge |
| Vehicle | vehicle, car, bus, school bus, truck, convertible, tank, patrol car, motorcycle, scooter, bicycle, train, streetcar, airplane, helicopter, battleplane, ship, boat, canoe, seat, driver seat, tire, steering wheel, car interior, driving |

Notes: The words were translated from Japanese into English. Although all original Japanese words were single, some consist of two English words after translation.

